# Membrane-Dependent Amyloid Aggregation of Human BAX α9 (173-192)

**DOI:** 10.1101/2020.10.20.347567

**Authors:** David A. Price, Tayler D. Hill, Kaitlyn A. Hutson, Blaze W. Rightnowar, Sean D. Moran

## Abstract

Mitochondrial outer membrane permeabilization, which is a critical step in apoptosis, is initiated upon transmembrane insertion of the C-terminal α-helix (α9) of the pro-apoptotic Bcl-2 family protein BAX. The isolated α9 fragment (residues 173-192) is also competent to disrupt model membranes, and the structures of its membrane-associated oligomers are of interest in understanding the potential roles of this sequence in apoptosis. Here, we used ultrafast two-dimensional infrared (2D IR) spectroscopy, thioflavin T binding, and transmission electron microscopy to show that the synthetic BAX α9 peptide (α9p) forms amyloid aggregates in solution and on the surfaces of anionic small unilamellar vesicles (SUVs). Its inherent amyloidogenicity was predicted by sequence analysis, and 2D IR spectra reveal that SUVs modulate the β-sheet structures of the resulting amyloid species. These results contradict prior models of transmembrane α9p pores and motivate further examination of the formation or suppression of BAX amyloids in apoptosis.

## 1. Introduction

The Bcl-2 family of proteins regulates the mitochondrial pathway of apoptosis,^1-3^ and dysfunction of one or more members can contribute to cancers,^4-6^ neurodegenerative diseases,^7,8^ and developmental disorders.^9,10^ The pro-apoptotic member BAX plays a central role in apoptosis via mitochondrial outer membrane permeabilization (MOMP), in which BAX oligomerizes on the mitochondrial outer membrane (MOM) and forms pores, which in turn release proteins such as cytochrome c and SMAC/DIABLO that promote downstream caspase activation and cell death.^3,11,12^ The early events in this process, including the insertion of the BAX C-terminal helix (α9) into the MOM and homodimerization, are increasingly well-characterized,^1,13-16^ but the molecular-level structure of the oligomeric BAX pore remains unknown. Additionally, non-MOMP functions of BAX have been reported^17^ and clusters containing hundreds to thousands of BAX monomers have been implicated in mitochondrial fission,^18,19^ suggesting multiple modes of action that likely involve undiscovered structural rearrangements.

In prior work, Tatulian and co-workers showed that the C-terminal fragment of human BAX, spanning α9 residues 173-192, is competent to permeabilize anionic liposomes^20^ and forms β-rich assemblies on supported lipid bilayers (SLBs).^21^ From these results, they proposed an octameric pore with a dimeter of 20-22 Å and mixed α/β conformation.^21^ Secondary structure information was derived primarily from analysis of FTIR spectra using Gaussian fitting protocols. Although such methods are useful for global characterization of peptide ensembles, they are prone to error from over-fitting^22^ and cannot easily distinguish between uniform structures and mixed states; for unambiguous assignment of spectra, specific attention must be paid to the underlying physical phenomena that influence vibrational frequencies and lineshapes. For example, low-frequency (∼1620 cm^−1^) amide I signals typically arise from excitonic coupling within extended β-sheets, such as those in amyloid aggregates,^23-27^ and differs from higher-frequency (≳1630 cm^−1^) features associated with β-sheets in native proteins.^28-30^ Similar features were observed in the previous data and cannot be accounted for by the small, two-stranded antiparallel β-sheets in the proposed BAX α9 pore model.^21^ Thus, alternative structures must be considered, and doing so may shed new light on the diverse roles of BAX and its regulation in cell survival and death.

In this study, we examined the aggregation of a synthetic human BAX α9 peptide (α9p) with the sequence H_2_N-VTIFVAGVLTASLTIWKKMG-CO_2_H. Using sequence-based structure analysis, we found that this region of the BAX sequence has a high β-aggregation and amyloid propensity. We then characterized α9p aggregates formed in the absence and presence of anionic small unilamellar vesicles (SUVs) using ultrafast two-dimensional infrared (2D IR) spectroscopy. This technique, which is described in detail elsewhere,^25,29,31-33^ provides substantial advantages over FTIR spectroscopy, including increased resolution of congested spectra, enhancement of vibrational modes with large transition dipole moments, and the ability to detect vibrational coupling through cross-peaks. Our 2D IR results, supported by thioflavin T (ThT) binding and transmission electron microscopy (TEM), show that α9p forms stable amyloid aggregates and that their formation and β-strand organizations are dependent on interactions with phospholipid bilayers.

## 2. Results

First, we used sequence-based structure analysis to determine β-aggregation or amyloid propensity of the full-length human BAX sequence (Figure 1). Four different algorithms (AGGRESCAN,^34^ TANGO,^35-37^ PASTA 2.0,^38^ and MetAmyl^39^) showed convergent predictions of high propensities localized within the C-terminal α-helix (α9) and some elevated propensities within α5 and α6, which are also known to associate with membranes.^16,40^ Here, the entire α9 region is predicted to form β-sheets, in contrast with the previous α/β pore model.^21^ Although these predictions are broadly consistent with the ability of α9p to form β-strands, none of these tools account for bilayer interactions, so independent aggregation trials in the absence and presence of model membranes is required to determine their effects.

**Figure 1.**
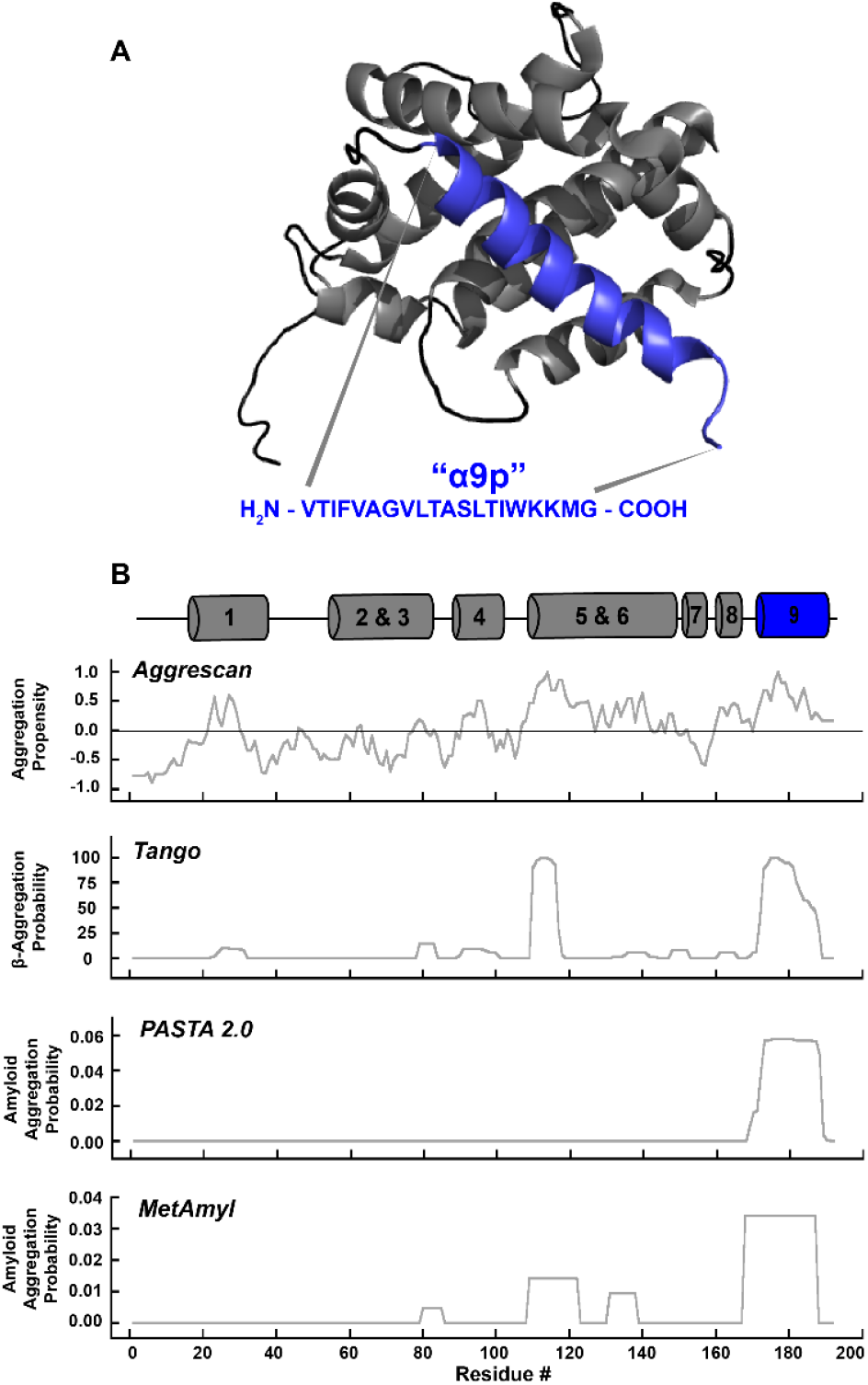
Structure and aggregation propensity of wild type human BAX. (A) Solution structure of BAX (PDB ID: 1F16)^41^ with helices α1-α8 in grey and α9 in blue, showing the sequence synthesized in this work (α9p). (B) Aggregation and amyloid propensities of full-length BAX protein via sequence analysis extracted from AGGRESCAN,^34^ TANGO,^35-37^ PASTA 2.0,^38^ and MetAmyl.^39^

To examine the aggregation of α9p in solution, we diluted 6 mM stocks of the disaggregated peptide to a final concentration of 150 μM into a D_2_O buffer containing 20 mM Tris (pD 7.5) and 100 mM NaCl, and followed the signal in the amide I region over time using rapid-scan 2D IR spectroscopy.^30,33,42^ Immediately after dilution, two diagonal peak pairs near ω_pump_ = 1645 cm^−1^ and ω_pump_ = 1675 cm^−1^ appeared, consistent with a largely disordered initial structure comprising random coil and β-turns, respectively (Figure 2A).^29,33,43-45^ Following a variable lag phase, a transition to β-sheets was observed by the growth of a broad diagonal peak pair between ω_pump_ = 1600 – 1635 cm^−1^ and concomitant reduction in the coil feature (Figure 2B). Eventually, a stationary phase is reached in which the spectrum is dominated by a narrow peak pair at ω_pump_ = 1620 cm^−1^ (Figure 2C). The low-frequencies of such signals arise from delocalized amide I normal modes oriented perpendicular to hydrogen-bonded β-strands, and the size of the resulting red shift scales with coupling constants that depend on the distances and angles between oscillators and follows an asymptotic trend with the number of strands in the β-sheets.^24,26,27,46,47^ Similar kinetic behavior observed via ThT fluorescence (Figure S1A), and TEM images (Figure S2A) of late-stage (24 h) aggregates confirm the formation of amyloid fibrils as was predicted from the sequence (Figure 1). Notably, the appearance and disappearance of the ∼1610 cm^−1^ diagonal signal (Figure 2D) at the onset of aggregation indicates the formation of a transient β-rich intermediate with different coupling parameters.^48^

**Figure 2.**
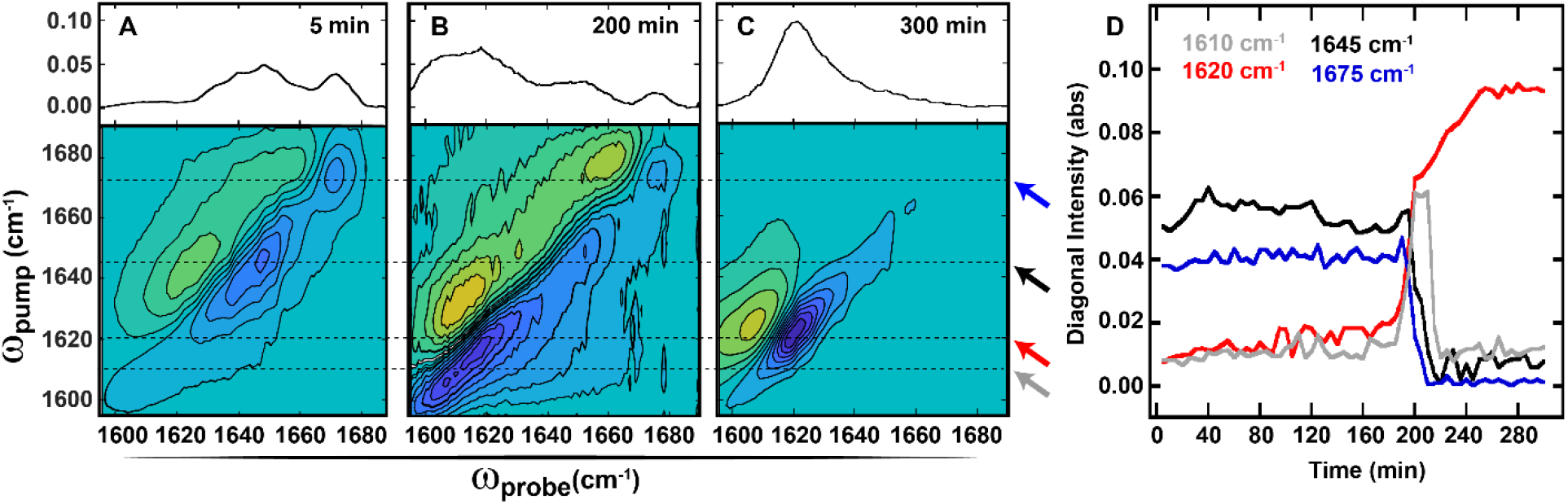
Aggregation behavior of α9p in aqueous buffer. (A-C) (Bottom) 2D IR spectra with (Top) corresponding diagonal slices for α9p in buffer at (A) early, (B) intermediate, and (C) late time points. Spectra are normalized to the largest amide I intensity in the stationary-phase (>250 min) spectrum and diagonal slices, which reflect the ν(0-1) bleach signals, are reversed in sign for comparison to FTIR absorbance. Horizontal lines are drawn through diagonal frequencies discussed in the main text and monitored in (D). (D) Diagonal maxima at 1610 cm^−1^ (grey), 1620 cm^−1^ (red), 1645 cm^−1^ (black), and 1675 (blue) cm^−1^ during α9p aggregation. Kinetic traces are color coded as indicated by arrows in A-C (*right*). Additional trials are shown in Figure S3.

Close inspection of the infrared spectra after 24 h incubation provides additional information about the orientation of β-strands in these aggregates. The FTIR spectrum and 2D IR diagonal slice (Figure 3A, top), are dominated by a ∼1620 cm^−1^ signal, and no other features are resolved above noise. However, in the 2D IR spectrum (Figure 3A, bottom) this feature (i) is accompanied by weak diagonal intensity in the coil region (ii) and near ω_pump_ = 1685 cm^−1^ (iii). Weak cross-peaks (iv) are also observed between the high- (iii) and low-frequency (i) features, indicating that these modes exist within the same coupled array. The high-frequency feature may arise from either β-turns or intra-strand coupling of amide I modes, and although these are difficult to distinguish in unlabeled samples,^47^ the low intensity is indicative of a likely parallel-stranded configuration (*vide infra*).^26,45,46^

**Figure 3.**
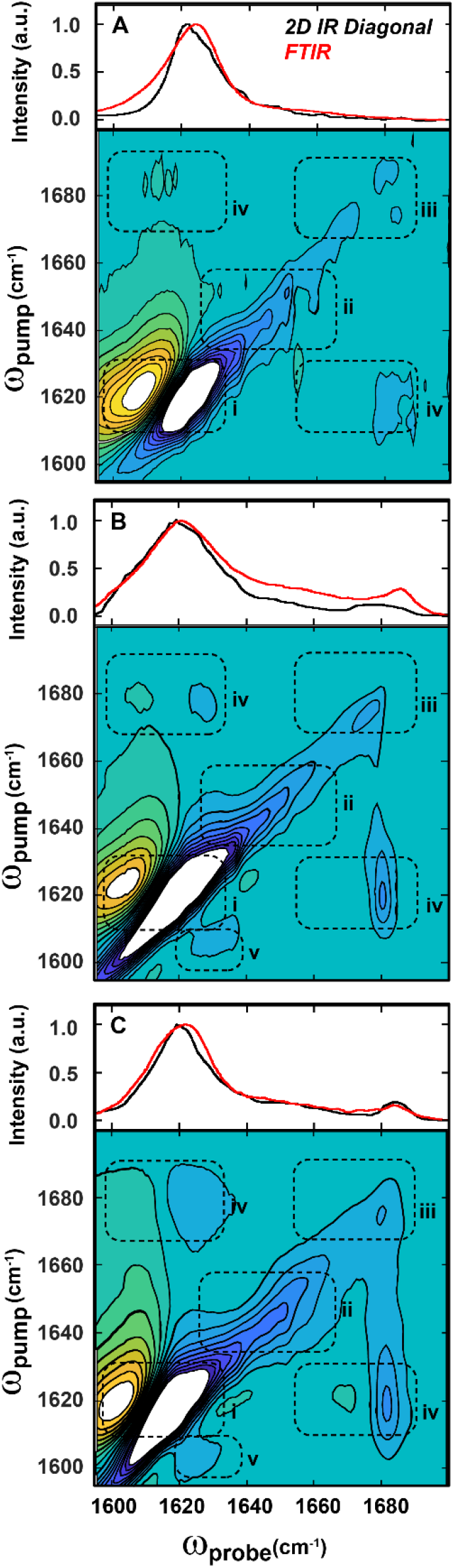
2D IR and FTIR spectra of mature (24 h) α9p aggregates. A-C: (Bottom) 2D IR spectra with (Top) corresponding diagonal slices (black line) and FTIR spectra (red line) for mature α9p aggregates alone in solution (A), and in the presence of 10 mM (B) and 1 μM (C) POPC:POPG SUVs. In all cases, α9p was incubated at room temperature for ∼24 h Tris/D_2_O buffer (pD = 7.5) to ensure aggregation. 2D IR spectra are scaled to 50% of their respective amide I maxima and peaks in regions i-iv correspond to structural features discussed in the text.

Next, we performed the aggregation experiments using suspensions of 3:1 POPC:POPG SUVs in the same buffer at a peptide:lipid ratio of 1:65 (10 mM total lipids). We observed rapid precipitation immediately following dilution; ThT fluorescence increased within ∼5 minutes (Figure S1B), and the resulting precipitates contained both SUVs and networks of fibril-like structures associated with their surfaces (Figure S2B). The accelerated formation of these species precluded 2D IR kinetics experiments, but in spectra collected after 24 h we observed a strong ∼1620 cm^−1^ signal that indicates amyloid formation (Figure 3B). In contrast to the aggregates formed in solution, this feature is broadened in both the FTIR spectrum and the 2D IR diagonal slice, and the high-frequency feature near 1685 cm^−1^ is clearly resolved (Figure 3B, top). These features (i, iii) are apparent along the diagonal in the 2D IR spectrum (Figure 3B, bottom) and the signal in the coil region (ii) remains relatively weak. The increase in intensity of the high-frequency feature, and enhancement of cross-peaks (iv) compared to Figure 3A, indicates that the α9p amyloids formed on SUVs have an alternative topology that includes a strongly coupled β-turn, antiparallel β-strands, or both.^23,24,46,47^ Additionally, the signal near 1610 cm^−1^ resembles low-frequency amide I modes in mature fibrils of Aβ, which were previously attributed to a sub-population of aggregates with enhanced inter-strand coupling.^48^ However, the presence of a cross-peak (v) to the dominant excitonic mode (i) indicates that different degrees of delocalization occur within the same structures, possibly as a result of structural inhomogeneity along β-strands within β-sheets. Interestingly, a similar low-frequency signal is observed in the intermediate states during lipid-free aggregation (Figure 2B,D) so it is possible that aggregation pathways diverge upon trapping of intermediates on SUVs followed by subsequent growth of aggregates with an altered topology.

Finally, we repeated the lipid-dependent aggregation experiments using a reduced concentration of SUVs (1 μM total lipids) for a peptide:lipid ratio of 150:1. Under these conditions, only a small fraction of α9p molecules can interact with bilayers, and a mixture of bilayer-dependent and bilayer-independent aggregates may be expected. Again, aggregation proceeded rapidly (Figure S1C) and aggregates containing SUVs and fibrillar structures were observed (Figure S2C). The FTIR and 2D IR spectra of the aggregates (Figure 3C) reproduced the features of those collected in the presence of excess lipids (Figure 2B), indicating that amyloid aggregation of α9p proceeds to completion; surprisingly, the intensities of the high-frequency (∼1685 cm^−1^) diagonal feature (iii) and its associated cross-peaks (iv) were further enhanced. Thus, the predominant structure resembles that of the amyloid aggregates formed with excess lipids (Figure 3B), and the contribution of bilayer-independent aggregation is minor. These results support a model of rapid nucleation of α9p on anionic bilayers followed by amyloid aggregation from a topologically distinct state that persists in a stable conformation for at least 24 hours.

## 3. Discussion

This study provides compelling evidence through sequence analysis, infrared spectroscopy, ThT fluorescence, and TEM imaging, that the C-terminal fragment (residues 173-192) of human BAX (α9p) forms amyloid fibrils in solution and on the surfaces of anionic (POPC:POPG) SUVs. The 2D IR data reveals that while these two classes of aggregates contain extended β-sheets that span the majority of the residues, they differ in the organization of their β-strands. Specifically, α9p aggregates formed in solution are parallel-stranded and those formed on SUVs have a more complex topology. In future studies, the detailed structures of these aggregates can be determined by combining 2D IR spectroscopy with isotope editing in order to reveal contacts between specific peptide bonds.^47,49-51^

By comparing our results to those of Tatulian and co-workers, we gain new insight into the structures of BAX α9p assemblies formed on model phospholipid bilayers. First, we consider the similarities between the two sets of results. Our FTIR and 2D IR spectra reproduce the strong signal at ∼1620 cm^−1^, which is well-known to arise from vibrational excitonic coupling in extended β-sheets^23,26,27,31,46^ and was likely mis-assigned to two-stranded structures in the α/β pore model.^21^ In the 2D IR spectra, we also resolve previously unassigned signals at ∼1610 cm^−1^ and ∼1685 cm^−1^, and cross-peaks show that they arise from coupled modes in the same assemblies. The latter high-frequency mode is consistent with the presence of antiparallel β-strands, albeit within amyloid aggregates instead of oligomeric transmembrane pores. The primary difference in our results is a reduced contribution of α-helices and disordered structures, which appear between 1635 – 1665 cm^−1^.^44^ It is possible that some differences in membrane composition, curvature, and stability could bias the system towards a stable transmembrane structure, but this does not account for the presence of the aforementioned excitonic signal in spectra of SLB-associated α9p.^21^ Instead, the pore model was more likely based on a mixture of states, including pre-amyloid and amyloid structures (cf. Figure 2A-C) that were unresolvable by FTIR. Although the final stable state of α9p is clearly amyloid in nature, this does not necessarily preclude the formation of transient pores along the bilayer-dependent aggregation pathway. Such kinetic details may become accessible via 2D IR with increased scan rates and fast mixing techniques.^52,53^

To date, structural studies of BAX have focused on the early events in MOMP, where transmembrane insertion of α9, homodimerization, and oligomerization are well-established.^1,11-16^ The results presented here show that the same region of sequence is highly amyloidogenic, and the resulting structures are modulated by model phospholipid bilayers. Clearly, more experiments will be required to establish whether the same phenomena occur in full-length BAX upon its interactions with mitochondrial membranes. However, the strong amyloid propensity of α9p suggests that the amyloid state of BAX is accessible under physiological conditions and must be considered in any detailed mechanistic model of apoptosis. Importantly, our observations generate new hypotheses regarding the possible functions of BAX. One possibility is that the amyloid pathway is suppressed *in vivo* by specific interactions of α9 with other regions of the BAX sequence, by structural preferences imposed by the MOM itself, or by other proteins involved in MOMP. Here, misfolding and amyloid aggregation of BAX would alter the balance of pro-apoptotic and anti-apoptotic Bcl-2 family proteins, perturbing pathways that regulate cell survival and death. Another possibility is that its amyloid-like structures (including intermediates) are directly involved in cell death as mitochondrial analogs of cytotoxic amyloids such as Aβ ^54-56^ and some antimicrobial peptides.^57,58^ Such structures could evolve from BAX pores,^11,12^ occur in late-stage BAX clusters,^18,19^ or form within non-canonical pathways of mitochondrial disruption and dysfunction.^17^ Finally, the recent observation that full-length BAX is incorporated into fibrils formed by the anti-apoptotic peptide humanin suggests that such heteroamyloids could act to suppress BAX function via sequestration in an inactive form.^59,60^ Thus, our results motivate the continued examination of the amyloid state of BAX *in vitro* and *in vivo* to further understand the molecular mechanisms of apoptosis.

## 4. Materials and Methods

### 4.1. Sequence analysis

The publicly available aggregation prediction servers AGGRESCAN (bioinf.uab.es/aggrescan),^34^ TANGO (tango.crg.es),^35-37^ PASTA 2.0 (protein.bio.unipd.it/pasta2),^38^ and MetAmyl (metamyl.genouest.org)^39^ were used to predict the aggregation propensities across the full-length human BAX protein (UniProt Accession ID: Q07812, PDB ID: 1F16, Figure S4). The AGGRESCAN server requires no input parameters outside of the amino acid sequence. For TANGO analysis, the option “no protection” was used for both the N- and C-termini. The pH, temperature, ionic strength, and concentration were set to 7.5, 298.15 K, 0.120 M, and 0.000150 M, respectively, to reflect solution conditions used for BAX α9 peptide experiments throughout this work. For PASTA 2.0 analysis, the Region (90% specificity) threshold was used which corresponds to top pairing energies and an energy threshold of 22 and < −2.8 PEU (PEU, 1.0 PASTA Energy Unit = 1.192 kcal/mol), respectively, which allow for high confidence in aggregation region detection. In the MetAmyl analysis, the “best global accuracy” threshold was applied for the BAX aggregation prediction.

### 4.2. Peptide synthesis and purification

The BAX α9 peptide (α9p) H_2_N-VTIFVAGVLTASLTIWKKMG-CO_2_H was synthesized in the solid state using standard fluorenylmethyloxycarbonyl (Fmoc) chemistry on Gly-Wang resin using an AAPPTec FOCUS XC automated peptide synthesizer. The product was purified by reverse-phase HPLC using a Hitachi LaChrom Elite system equipped with a Vydac semi-preparative C4 column. Acetonitrile and water, supplemented with 0.1% 2,2,2-trifluoroacetic acid (TFA) and 10% 2,2,2-trifluoroethanol (TFE), were used as the mobile phase for HPLC purification, with a linear gradient of 30% to 37% acetonitrile:TFE:TFA over 60 min. The pure peptide was confirmed using a Bruker Daltonic MALDI-TOF Mass Spectrometer (Figure S5).

### 4.3. Sample preparation

Purified α9p was resuspended in deuterated hexafluoroisopropanol (d-HFIP) to deuterate exchangeable sites and promote disaggregation of peptides.^27^ Concentrated (6 mM) peptide/d-HFIP solution was diluted into Tris-buffered (20 mM Tris, 100 mM NaCl, pD/pH = 7.5) D_2_O or H_2_O, either with or without lipid vesicles, to a final peptide concentration of 150 μM and incubated at room temperature for ∼ 24 h to ensure aggregation unless otherwise noted. For SUV preparation, chloroform solutions of 1-palmitoyl-2-oleoyl-sn-glycero-3-phosphocholine (POPC) and 1-palmitoyl-2-oleoyl-sn-glycero-3-phosphoglycerol (POPG) (Avanti Polar Lipids, Alabaster, AL) were combined in a ratio of 3:1 POPC:POPG and dried under vacuum. Lipid films were resuspended to 10 mM lipid concentration in the appropriate buffer and mixed vigorously for 1 h. SUVs were prepared by sonicating the resulting suspensions for 10 min.^61^ For experiments at 10 mM lipids, SUV suspensions were used as prepared and for experiments at low lipid concentrations SUV suspensions were diluted into identical buffers to the desired concentration.

### 4.4. Infrared spectroscopy

Sample and background FTIR spectra were collected with 128 scans at room temperature using a Jasco 6800 FTIR spectrometer equipped with a PIKE Technologies MIRacle ATR Accessory and DTGS detector with a resolution of 1 cm^−1^. Buffer backgrounds were subtracted and residual baseline correction was performed in MATLAB using low-order polynomials. For 2D IR spectroscopy, ∼17 μJ mid-IR pulses with (∼100 fs) were directed into a 2DQuick Array spectrometer (Phasetech Spectroscopy, Inc., Madison, WI) equipped with a 128 x 128 pixel MCT array detector as previously described.^62^ All spectra were collected with perpendicular pump and probe polarizations and four-frame phase cycling to reduce interference from pump scatter. Probe frequencies were calibrated to the absorbances of 4-nitrobenzaldehyde in dichloromethane (1605, 1676, and 1709 cm^−1^), used as an external standard. All instruments were purged continuously with dry air to minimize contributions from water vapor. 2D IR spectra were processed in MATLAB using low-order polynomial background correction and Hamming apodization.^49^

### 4.5. Thioflavin T Fluorescence

For Thioflavin T experiments, concentrated peptide/d-HFIP solutions were diluted directly into a quartz cuvette containing Tris-buffer, ThT (25 μM) and SUVs. Emission spectra between 450 – 600 nm were collected in 30 s intervals using a Cary Eclipse spectrofluorometer (λ_ex_ = 440 nm), and emission intensities at 480 nm were monitored over time.

### 4.6. Transmission Electron Microscopy

For TEM experiments, 5 µL of buffered α9p samples, with or without SUVs, were deposited onto formvar coated nickel grids (Electron Microscopy Sciences, Hatfield, PA) and allowed to incubate for 5 min at room temperature. Excess buffer was removed by blotting, and the grids were dried in air for 10 min and stained with 2% Uranyl Acetate. TEM images were collected with a Hitachi H-7650 Transmission Electron Microscope using an acceleration voltage of 60 kV.

## Acknowledgements

The authors wish to thank the Southern Illinois University IMAGE center for assistance in the collection of the TEM images. The authors wish to thank the SIU Mass Spectrometry Facility director Dr. Mary Kinsel for training and assistance pertaining to the Bruker Daltonic MALDI TOFMS results included in this publication.

## Notes

The authors declare no competing financial interest.

