## Supplemental Figures S1-S5 for "Membrane-Dependent Amyloid Aggregation of Human BAX α9 (173-192)"

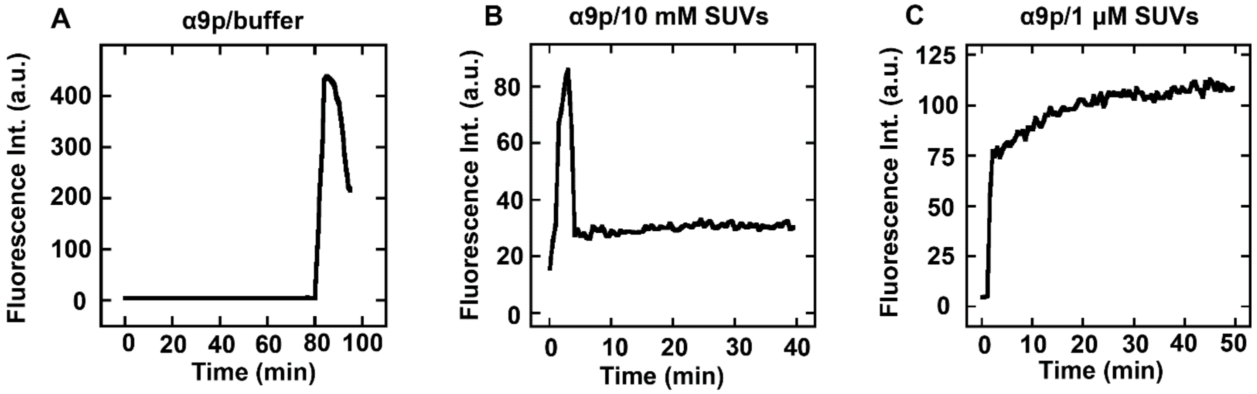


**Figure S1.** Thioflavin T (ThT) fluorescence analyses of BAX α9p aggregation. ThT fluorescence ($\lambda_{ex}$ = 440 nm) was measured at 480 nm over time during the aggregation of 150 μM α9p in the absence and presence of lipid vesicles (A) ThT fluorescence during α9p aggregation in buffer (pH = 7.5) showing a ~80 min lag phase followed by a rapid increase in fluorescence during the growth phase and subsequent decrease in fluorescence upon precipitation of ThT/α9p aggregates from solution. (B) ThT fluorescence during α9p aggregation in the presence of 10 mM and POPC:POPG (3:1) SUVs. Fluorescence increases rapidly within 5 minutes and decreases before 10 min upon the precipitation of ThT/α9p vesicle associated aggregates. (C) ThT fluorescence during α9p aggregation in the presence of 1 μM and POPC:POPG (3:1) SUVs. Fluorescence increases rapidly within 5 minutes upon aggregation of α9p and slowly increases to a plateau near 20 min.

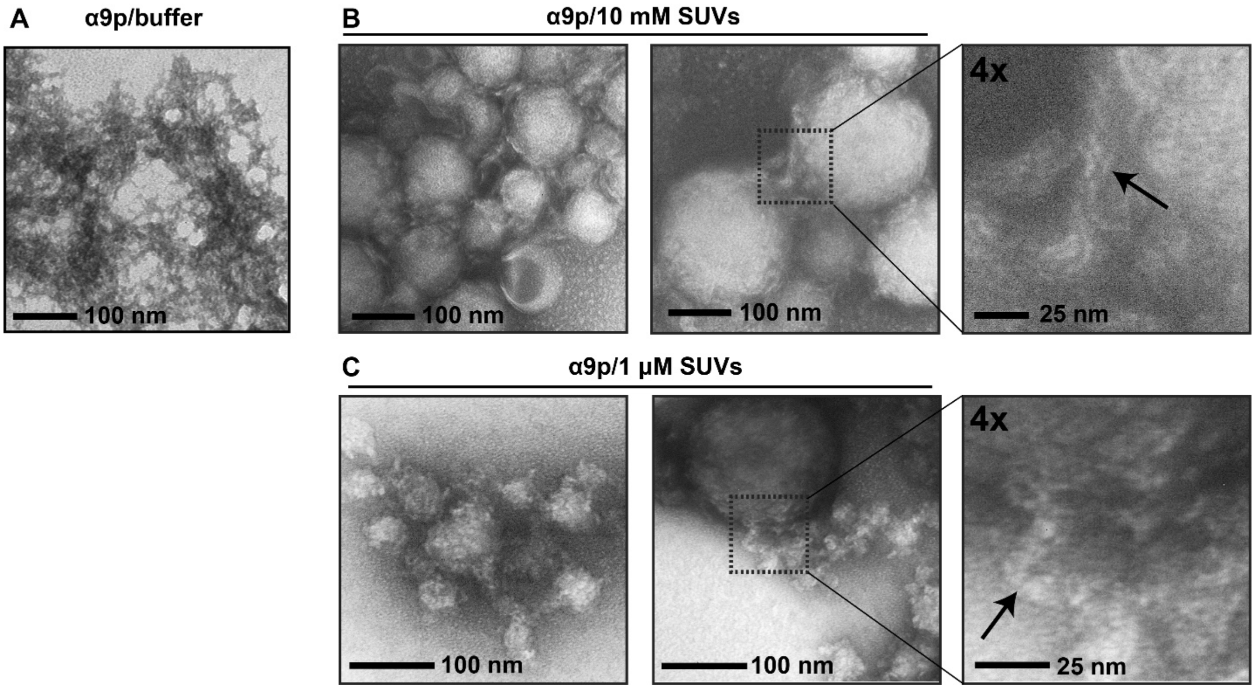


**Figure S2.** TEM images of mature α9p aggregates. (A) TEM image of mature α9p aggregates showing web-like fibrillar structure. (B) TEM images of α9p/SUV cluster aggregates formed from α9p in the presence of SUVs (10 mM total lipids). *Left/Middle:* TEM images of α9p/SUV cluster aggregates showing linkage of vesicles by fibrillar aggregates. The dashed box in the middle panel shows region of image magnified in the right panel. *Right:* Detail of TEM image (4x zoom) with black arrow pointing to aggregates associated with the vesicle surface. (C) TEM images of α9p/SUV aggregates formed from α9p in the presence of SUVs (1 µM total lipids). *Left/Middle:* TEM images of α9p/SUV aggregates showing fibrillar aggregates protruding from vesicle surface. The dashed box in the middle panel shows region of image magnified in the right panel. *Right:* Detail of the TEM image (4x zoom) with black arrow pointing to aggregates.

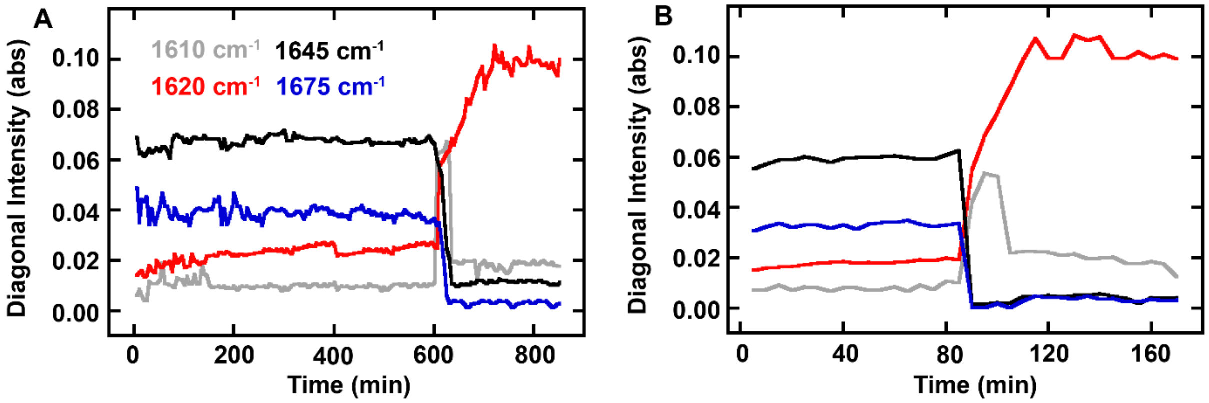


**Figure S3.** BAX α9p aggregation kinetics in absence of SUVs. 2D IR diagonal maxima at 1610 $cm^{-1}$ (grey), 1620 $cm^{-1}$ (red), 1645 $cm^{-1}$ (black), and 1675 (blue) $cm^{-1}$ during α9p aggregation for 2 separate trials.

```
                                                                 60
MDGSGEQPRG GGPTSSEQIM KTGALLLQGF IQDRAGRMGG EAPELALDPV PQDASTKKLS
                                                                120
ECLKRIGDEL DSNMELQRMI AAVDTDSPRE VFFRVAADMF SDGNFNWGRV VALFYFASKL
                                                                180
VLKALCTKVP ELIRTIMGWT LDFLRERLLG WIQDQGGWDG LLSYFGTPTW QTVTIFVAGV
       190
LTASLTIWKK MG
```

**Figure S4.** Amino acid sequence of the wild-type human BAX protein (UniProt Accession ID: Q07812, PDB ID: 1F16). The sequence of BAX α9p synthesized in this work (residues 173-192) is underlined.

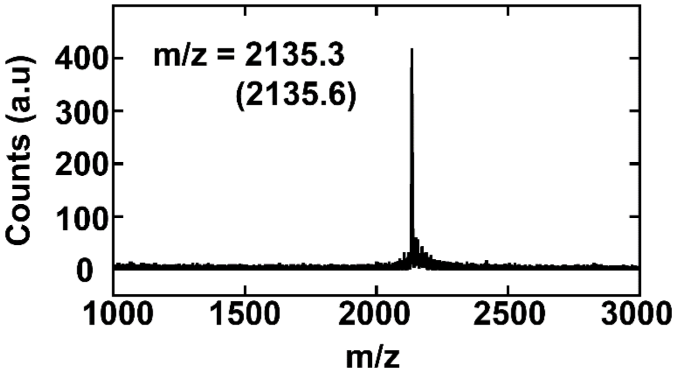


**Figure S5.** MALDI-TOF mass spectrum of synthetic α9p. The observed m/z is shown in panel with expected value in parentheses.
